# Upscaling of *In Vivo* HR-pQCT Images Enables Accurate Simulations of Human Microstructural Bone Adaptation

**DOI:** 10.1101/2020.05.13.093328

**Authors:** Nicholas Ohs, Duncan C. Tourolle né Betts, Penny R. Atkins, Stephanie Sebastian, Bert van Rietbergen, Michael Blauth, Patrik Christen, Ralph Müller

## Abstract

*In silico* trials of treatments in a virtual physiological human (VPH) would revolutionize research in the biomedical field. Hallmarks of bone disease and treatments can already be simulated in pre-clinical models and in *ex vivo* data of humans using microstructural bone adaptation simulations. The increasing availability of *in vivo* high resolution peripheral quantitative computed tomography (HR-pQCT) images provides novel opportunities to validate and ultimately utilize microstructural bone adaptation simulations to improve our understanding of bone diseases and move towards *in silico* VPH decision support systems for clinicians.

In the present study, we investigated if microstructural bone adaptation simulations of *in vivo* human HR-pQCT images yielded accurate results. Since high-resolution ground truth images cannot be obtained *in vivo*, we applied an *ex vivo* approach to study resolution dependence and the effect of upscaling on morphometric accuracy. To address simulation initialisation issues, we developed an input regularisation approach to reduce initialisation shocks observed in microstructural bone adaptation simulations and evaluated upscaling as a way to improve the accuracy of model inputs. Finally, we compared our *ex vivo* results to simulations run on *in vivo* images to investigate whether *in vivo* image artefacts further affect simulation outcomes.

## Introduction

Simulations are considered the third pillar of modern science next to models and experiments. In the biomedical field the creation of a virtual physiological human (VPH) is seen as one of the most important goals (Fenner et al. 2008). The vision of the VPH is to provide researchers with a model that allows rapid hypothesis testing via *in silico* trials and provides doctors with a virtual patient as a decision-support system for their daily work (Viceconti and Hunter 2016). The role of bone, both structurally and physiologically, indicates that a validated model for microstructural bone adaptation and (re)modelling is a significant component to any VPH model. Previous studies have shown that various aspects of bone diseases and their treatments can be simulated in pre-clinical models (Müller 2005; Ruimerman et al. 2005; Schulte et al. 2013; Levchuk et al. 2014). Importantly, the advection based model by Adachi et al. (2001) can produce results comparable to population data when *ex vivo* images are used as an input (Badilatti et al. 2016). However, the translation of microstructural simulations to clinical image data has largely been constrained by the availability of high quality images and validation data.

With the introduction and increased use of HR-pQCT, large amounts of clinically relevant data have been gathered which provide the basis to validate and parameterise *in silico* models of bone (Boutroy et al. 2005, 2008; Sornay-Rendu et al. 2007, 2017; Kirmani et al. 2009; Nicks et al. 2012; Nishiyama et al. 2013; Nishiyama and Shane 2013; Yu et al. 2014; Zhu et al. 2014). However, combining current microstructural bone adaptation simulations with HR-pQCT is non-trivial. Existing simulations either utilize synthetic images (Ruimerman et al. 2005) or high-resolution micro-CT images which cannot be obtained clinically (Müller 2005; Schulte et al. 2013; Levchuk et al. 2014; Badilatti et al. 2016). Furthermore, HR-pQCT images tend to have more noise (Rajapakse et al. 2009) and other potential imaging artefacts, such as those due to movement (Pialat et al. 2012).

The reduction in resolution is a known obstacle for the translation of computational techniques from the lab into the clinical setting (Ţjong et al. 2012; Manske et al. 2015; Christen et al. 2016).

Alsayednoor et al. (2018), for example, showed that applying a single threshold to HR-pQCT images cannot yield both correct morphometric indices and mechanical properties on HR-pQCT images. However, *in silico* microstructural bone adaptations rely on having correct digital representations of morphometrics *and* mechanics as the simulation couples these two properties. To study the effects of image resolution in detail, convergence studies have been performed which showed a clear dependence on image resolution for various mechanical and morphometric parameters (Müller et al. 1996; Christen et al. 2016). In order to counteract this algorithmic dependence on image resolution, upscaling low-resolution images to desktop micro-CT images on which the algorithms have been validated can help to produce accurate results (Rajapakse et al. 2009; Schulte et al. 2019). For example, upscaling of magnetic resonance imaging (MRI) data has been shown to produce micro-FE results in good agreement with those from micro-CT images of a higher resolution (Rajapakse et al. 2009). While it is clear that upscaling does not yield the same effects as scanning at a higher resolution (Cooper et al. 2007), as upscaling cannot compensate for information missing in the image, techniques, like mesh refinement, are widely used in numerical applications to improve simulation accuracy by providing a better digital representation of the information contained in the images.

In previous studies using the algorithm by Adachi et al. (2001), initial iterations, which showed aberrant results, were regarded as part of the model initialization and excluded from analysis (Schulte et al. 2013; Badilatti et al. 2016). However, the exclusion of the initial iterations would lead to an inherent divergence between the clinical *in vivo* and *in silico* baseline models, which thus precludes direct comparison of simulation outcomes against clinically observed changes in bone microstructure, and hinders validation of the *in silico* model. The aberrant results of the initial iterations are caused by an initialisation shock, which is common when modelling coupled systems, like that of advection and finite-element methods by Adachi et al. (2001), and are related to mismatches between experimental input data and simulation parameters (Balmaseda and Anderson 2009; Mulholland et al. 2015). In the context of microstructural bone adaptation simulations, the results of these mismatches can be observed in the large sudden changes in parameters, such as the total bone volume or the overall structural stiffness. Reducing this shock behaviour allows for the inclusion of all simulation iterations, such that identical baseline models can be used for the clinical and *in silico* models and results can be directly compared.

The goal of the present study was to determine if microstructural simulations of *in vivo* human HR-pQCT images based on the approach by Adachi et al. (2001) yield accurate results, such that they could be used as part of a VPH. Since high-resolution ground truth micro-CT images cannot be obtained *in vivo*, we applied an *ex vivo* approach to study the resolution dependence and the effects of upscaling on morphometric accuracy over the course of a microstructural bone adaptation simulation. However, we first had to address two initialisation issues: initialisation shocks and disagreement of mechanics and morphometrics in model inputs at HR-pQCT resolution. Thus, we developed an input regularisation approach to reduce initialisation shocks observed in microstructural bone adaptation simulations and studied upscaling as a method to improve the accuracy of model inputs with respect to both mechanics and morphometry. Finally, we compared our *in silico* results to simulations run on *in vivo* images to investigate whether additional *in vivo* image artefacts affect simulation outcomes.

## Materials

### High Resolution *Ex Vivo* Micro-CT Images

Five distal radii were obtained from female cadavers at the Amsterdam Medical Center as part of a previous study (De Jong et al. 2016). The donors’ ages varied between 58 and 95 years and the bone volume fraction (BV/TV) of the samples varied from 7 to 20% where BV/TV was inversely related with age. The medical history of the cadaveric specimens was unknown. High resolution CT images were obtained at an isotropic voxel-size of 25 μm with a vivaCT 80 (70 kV, 114 μA, 300 ms integration time), a micro-CT device by Scanco Medical AG (Switzerland). Images were Gauss-filtered (sigma = 1.2, support = 1) and the trabecular region was hand-contoured by a trained operator for each scan using the software of the scanner manufacturer.

### *In Vivo* HR-pQCT Images

Five patients (four female, one male) were recruited at Innsbruck Medical University as part of a radius fracture study. Patients provided informed consent and participated in a study approved by the ethics committee of the Medical University of Innsbruck. The age of the patients ranged from 26 to 80 years and their BV/TV from 21 to 7%. In this study, images of the unfractured, contralateral radius were used. Scans were performed with an isotropic voxel-size of 61 μm using an XtremeCT II (68 kV, 1470 μA, 43 ms integration time), a clinical HR-pQCT device by Scanco Medical AG (Switzerland). Images were Gauss-filtered (sigma = 1.2, support = 1) and the trabecular region was hand-contoured by a trained operator using the software of the scanner manufacturer.

### Generation of Low-Resolution Images

The high-resolution *ex vivo* micro-CT grey-scale images were downscaled to resolutions of 40, 61, and 82 μm. These three resolutions will be referred to as low-resolutions in this paper. Currently, 61 and 82 μm are the highest resolutions available for clinical CT scanners (XtremeCT I and II, Scanco Medical AG, Switzerland), these resolutions will be referred to as the clinically relevant resolutions. Resizing was performed using the scikit-image (Van Der Walt et al. 2014) rescale function in Python (Python Software Foundation 2020) with third-order interpolation and anti-aliasing enabled. The binary hand-contoured masks for the high-resolution images were converted to a floating-point data-type and resized like the micro-CT images. Finally, a threshold of 50% of the maximum image value was applied to obtain binary masks for the low-resolution images.

### Generation of Upscaled Images

Upscaled images were created from the low-resolution images by applying the scikit-image resize Python function, again with third-order interpolation and anti-aliasing enabled, to a resolution of 25 μm. When creating the upscaled images, the conversion of image dimensions between the different resolutions was not unique (i.e. due to rounding of the integer image dimensions after scaling with a floating point number, differences in size of one voxel could occur). To ensure that the upscaled images have the exact same dimensions as the original images, the scikit image resize function was used. The resize function is identical to the rescale function with the exception that it resizes images to a target image dimension instead of resizing using a target scaling factor. For the upscaled images, the same hand-contoured masks for the trabecular region were used as for the high-resolution *ex vivo* micro-CT grey-scale images to avoid influences of differences in masks on our results. Herein, the images upscaled from 40, 61, and 82 μm to 25 μm are referred to as u40 μm, u61 μm, and u82 μm, respectively.

The 61 μm clinical *in vivo* HR-pQCT images were re-sampled using the scikit-image rescale function, as was used for the *ex vivo* micro-CT images, to generate datasets at resolutions of 25, 40, and 82 μm. The masks of the HR-pQCT images were similarly rescaled.

## Methods

### Remodelling Simulations

#### Micro-FE Analysis

For the micro-FE analysis, we used the parallel octree solver parOsol (Flaig 2012) on the supercomputer Piz Daint at the Swiss National Supercomputing Centre (CSCS, Lugano, Switzerland). Output parameters of strain energy density (SED) and the apparent compressive stiffness along the longitudinal axis were evaluated. Boundary conditions were determined using a load estimation algorithm developed by Christen et al. (2013). This algorithm tries to linearly combine three different load cases to achieve the most homogeneous SED distribution possible across the given bone structure. The target mean SED value was 0.02 MPa, as has been used previously (Christen et al. 2016). Furthermore, soft pads were added to the distal and proximal ends of the images with a pad-thickness of 246 μm and a Young’s modulus of 15 MPa, which has previously been found to improve the load estimation (Christen et al. 2013). For all experiments, we computed the load-estimation using the high-resolution files and applied the same loading conditions to the low-resolution and upscaled files. This method of load-estimation removes the voxel-size dependency of the algorithm as a confounding factor.

The micro-FE simulations for images with resolutions higher than 50 μm were run on a 50 μm hexahedral mesh since the mechanical signalling implemented in the microstructural bone adaptation simulation is roughly equivalent to a blurring with a sigma of 100 μm. Therefore, the additional resolution in the SED would not yield differing results. The use of the 50 μm hexahedral mesh also reduced the computational resources required to run simulations (i.e. for the 25 μm images, this reduction was an order of magnitude).

#### Remodelling Algorithm

The strain-adaptive *in silico* microstructural bone adaptation simulation by Adachi et al. (2001) was re-implemented in Python using NumPy (Van Der Walt et al. 2011) and pybind11 (Jakob et al. 2017). In short, this algorithm is iterative; for each step, SED (a result of the micro-FE simulation) is translated, via a mechanostat, into the velocity field of an adapted advection equation. Within this advection equation, the mass transfer is constrained within a proximity of the bone surfaces, which results in changes to the bone microstructure. Due to the implementation, all changes are limited to the trabecular region of the simulated structure.

#### Binary Model Generation from Micro-CT Data

The high-resolution images were segmented using a threshold of 450 mg HA/cm^3^, which has also been chosen in previous studies on this dataset (Müller et al. 1996; Christen et al. 2016), and the bone volume over total volume (BV/TV) for the trabecular regions was computed for reference. Finally, voxels identified as bone were set to 750 mg HA / cm^3^ and background voxels were set to 0 mg HA / cm^3^.

#### Regularized Model Generation

To ensure that the remodelling simulation operates only on the bone surface, the input to the algorithm is required to be binary except for surface voxels that can be represented with intermediate values. To compare the effects of using a conventional binary input or using an input allowing partially filled voxels at the surface layer, we implemented a regularization method that preserves information of the grey-scale image at the surface of bone structures (Figure 1, left). First, a regularization threshold was applied to each high-resolution grey-scale image. Then, surface voxels (empty voxels in direct face-to-face contact with full voxels) of the intermediate binary structure were identified using a Von Neumann neighbourhood. For each surface voxel, grey-scale values from the original grey scale image were converted to a value in the range of zero to one relative to the regularization threshold. The regularization threshold was chosen such that the grey-scale BV/TV of the resulting structure was identical to the one computed for the respective conventional binary structure. Finally, the entire structure was multiplied by the same density value as the conventional binary input (750 mg HA / cm^3^).

**Fig 1.**
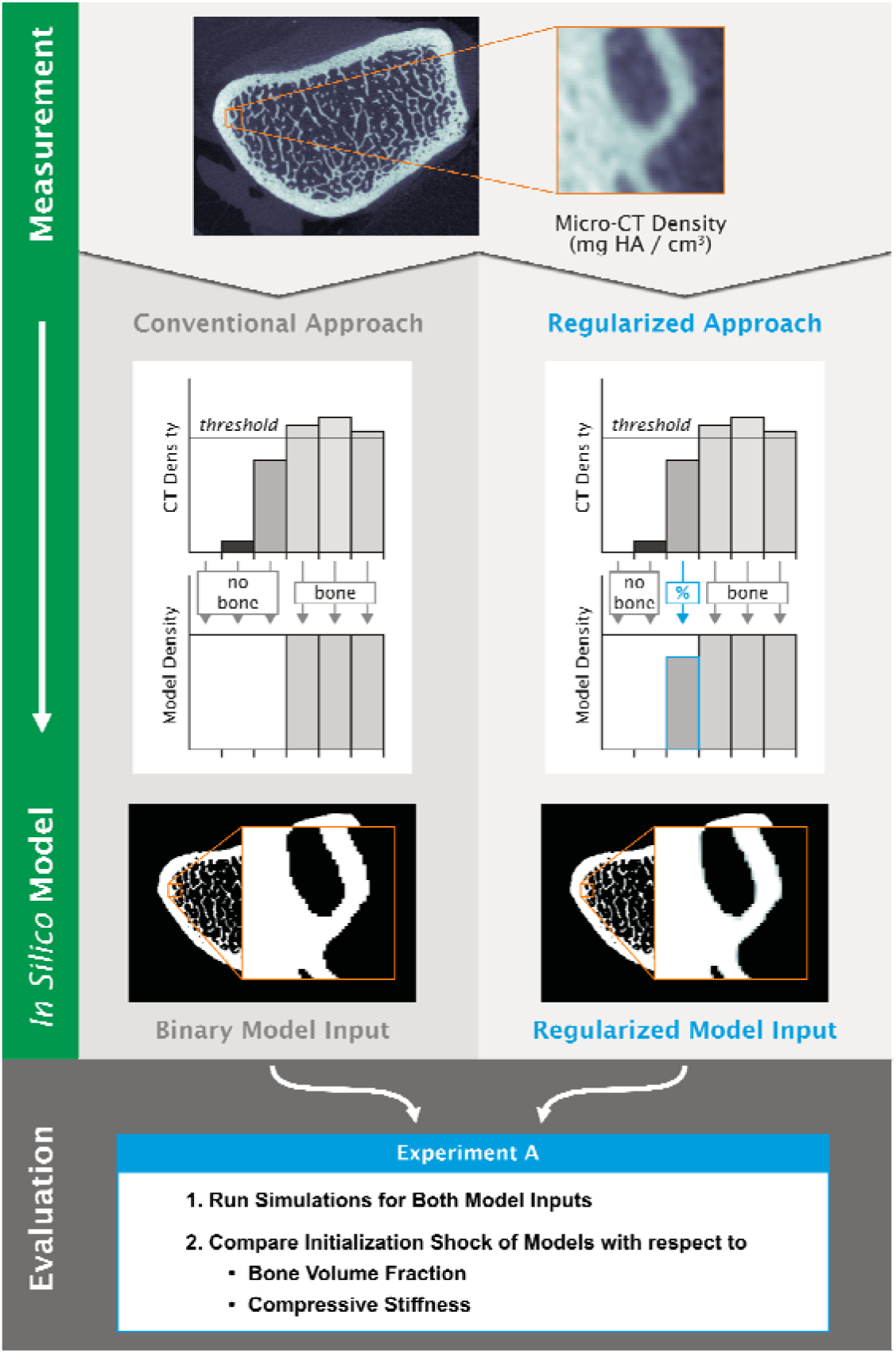
Experiment A (Effect of Regularized Input Models on Initialisation Shock) evaluated the reduction of initialisation shock behaviour in bone volume fraction and compressive stiffness due to a novel input regularization approach compared to the conventional input approach. Left: Illustration of the conventional threshold approach used to convert a CT image into a valid input for the load adaptation simulation. Right: Regularization approach

#### Parameters for Microstructural Bone Adaptation Simulation

Simulation parameters were chosen such that both formation and resorption were observed in the simulations and were used for all samples and resolutions. For this reason, a narrow lazy zone (0.0196 MPa to 0.0204 MPa) was chosen. The maximum velocity of the mechanostat was set to an arbitrary value of 12 μm/month. The slopes of the mechanostat were set to 8000 μm/year/MPa. The chosen value for slope resulted in generally high velocities and greater changes per time unit, due to the very narrow linear regime of the mechanostat. The choice of simulation parameters allowed for large differences between the different resolutions.

To ensure that the choice of time step between consecutive micro-FE calls did not alter the results, a time step of approximately 1.9 months was chosen. The simulated time period was set to 5 years, resulting in a shorter final iteration step.

### Study Design

The validity of using *in vivo* HR-pQCT data as an input for advection based microstructural bone adaptation simulations was investigated using four virtual experiments. Experiment A addresses the issue of initialisation shocks and compares the current approach of generating simulation input models with a novel regularization method. The regularized approach retains grey-scale information, allowing the simulation to initialise with a structure closer to the original one. The goal was to compare the behaviour of the two approaches during the initial iteration steps to identify the approach that exhibits the least amount of initialization shock. Experiment B compared the mechanical and morphological properties of regularized input models generated from high-resolution images, which had been downscaled or down- and then upscaled, that were bone volume fraction matched to the original high-resolution image. The aim of this experiment was to quantify differences between regularized input models with respect to mechanics and morphometrics that may confound simulations. In experiment C, microstructural bone adaptation simulations were run on models of all three *ex vivo* image sets (high-resolution, downscaled, and down-then upscaled) to assess if observed differences in the simulations were due to a lack of fidelity in the input data or the numerical grid. Finally, in experiment D, the *in vivo* images were rescaled to the same resolutions used in experiment C and the convergence was quantified with respect to resolution. The results were compared to those from experiment C to assess what effect differences in image quality and factors other than resolution have on the outcome of the simulation.

#### Experiment A: Effect of Regularized Input Models on Initialisation Shock

Two simulation input model generation approaches were compared (Figure 1, right), the current state of the art through binary representation and a regularized model with partially filled voxels at the surface. Both models were generated from the high resolution *ex vivo* micro-CT image data set. The partially filled voxels approximate a bone surface with sub-voxel precision. Microstructural bone adaptation simulations were run and the discontinuity in BV/TV and compressive stiffness for the initial simulation steps were quantified for both methods.

#### Experiment B: Effect of Upscaling on Mechanical and Morphometric Accuracy

For experiments B, regularized input models were produced for all three micro-CT image sets: high-resolution, low-resolution, and upscaled low-resolution images. In the following, these are called reference, low-resolution, and upscaled regularized input models. For all images, the reference BV/TV was always the one obtained from the respective binary high-resolution image.

To quantify the agreement in mechanical properties between reference, low-resolution, and upscaled regularized input models of the same bone structure, micro-FE analyses were performed on the data set at each of these resolutions, respectively (Figure 2). SED distributions, mean SED, mean static parameters (BV/TV, trabecular number (Tb.N), trabecular thickness (Tb.Th), trabecular spacing (Tb.Sp), and structural model index (SMI)), and standard deviations for all mean values were computed for comparison between the three regularized input model types. Additionally, the Kolmogorov-Smirnov statistic was computed between the SED of each sample of the low-resolution and upscaled regularized input models and the corresponding reference regularized input model to quantify mechanical agreement. Finally, the adequacy of the chosen threshold was assessed through comparison of the SED distribution from FE analysis of the low-resolution regularized input models for a range of thresholds (575 - 775 mg HA/cm^3^). The magnitudes of the peak SED were compared to the reference SED distribution.

**Fig 2.**
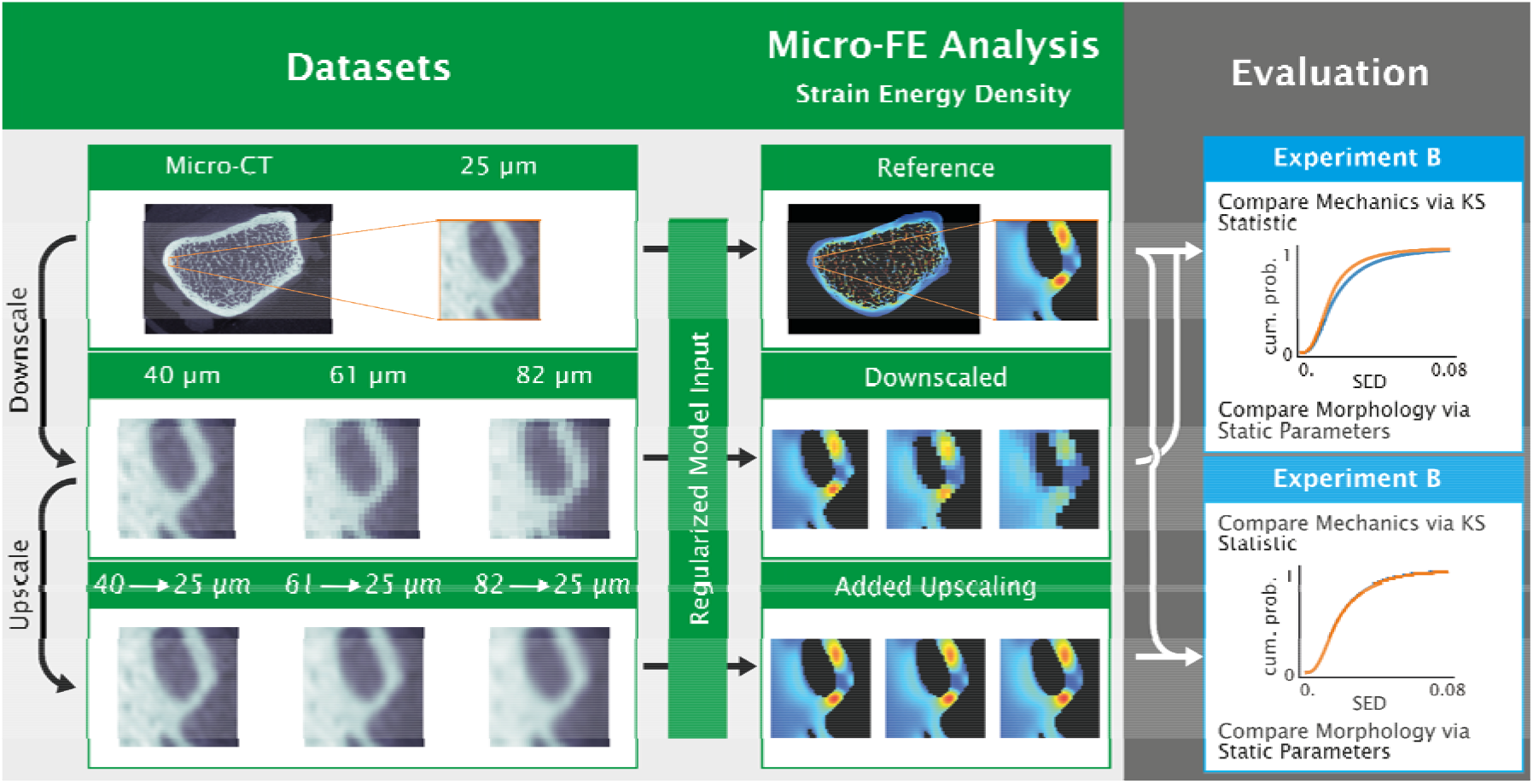
Experiment B (Effect of Upscaling on Mechanical and Morphometric Accuracy) evaluated the mechanical agreement between the different resolutions for matched BV/TV. For each micro-CT image, high-, low-resolution, and upscaled images were created (left column). Low-resolution: 40 μm, 61 μm, and 82 μm and upscaled images: Same downscaled resolutions as the low-resolution image but upscaled back to 25 μm. Micro-finite-element (micro-FE) models were generated using the regularization model generation approach of experiment A (Figure 1) and matching bone volume fraction (BV/TV) for each image to the reference (25 μm). A micro-FE analysis was run and strain energy density (SED) (shown in the jet colour-map) and static parameters were computed for each model. SED distributions between low and upscaled resolution images were compared using the Kolmogorov-Smirnov statistic to see which one more closely matches the reference high-resolution regularized input model mechanically. Mean static parameters were compared to see which one more closely matches the reference model morphologically

#### Experiment C: Effect of Upscaling on Morphometric Accuracy Throughout a Microstructural Bone Adaptation Simulation

Microstructural bone adaptation simulations were run on the input models generated for experiment B. The simulations run on the reference regularized input models were considered the best approximations of the *in vivo* remodelling process and were used as reference to quantify errors. These simulations are referred to as the reference simulations (Figure 3). Static parameters (BV/TV, Tb.N, Tb.Th, Tb.Sp, and SMI) and formed and resorbed volume over time were computed.

**Fig 3.**
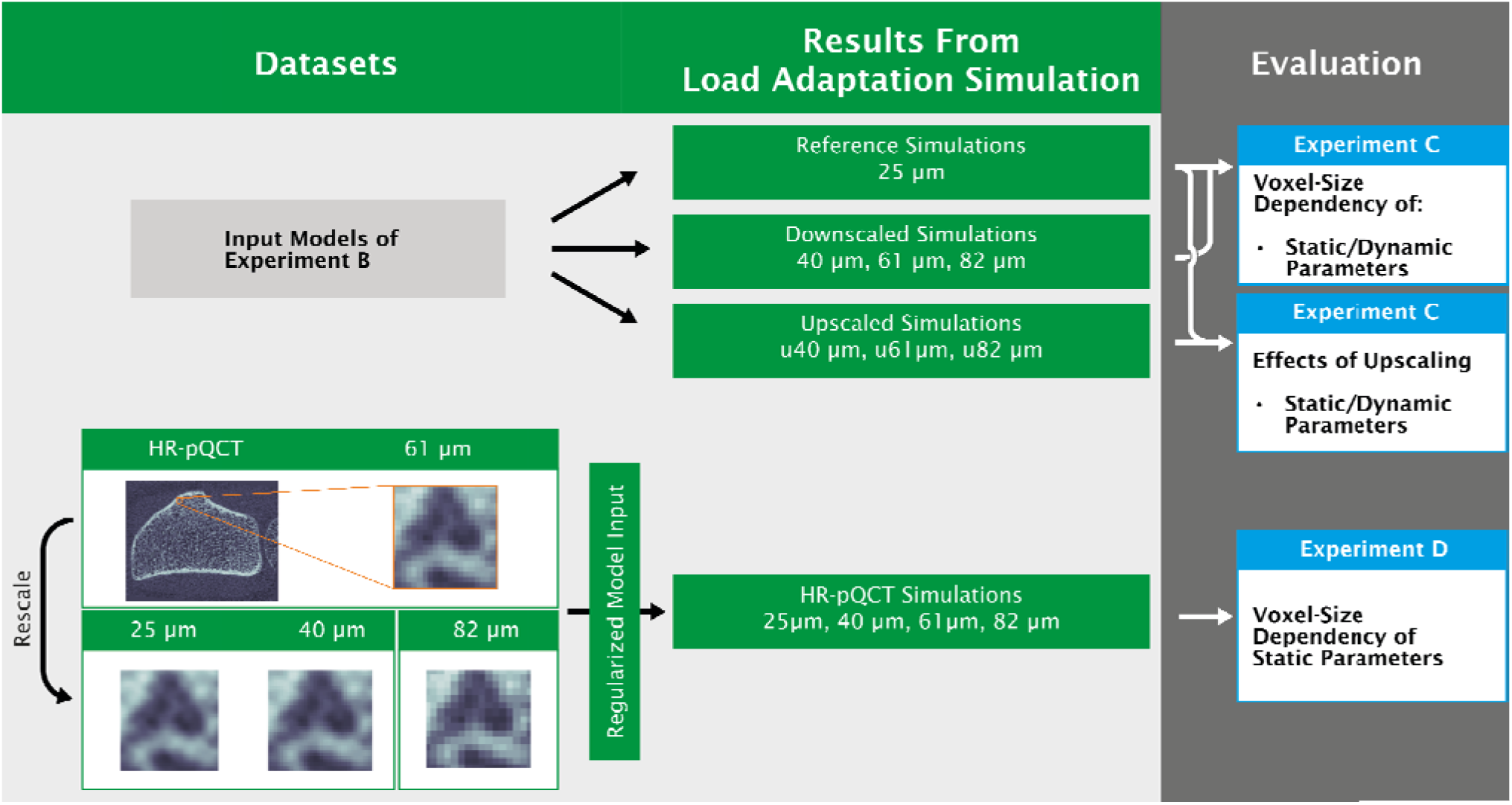
Overview of experiments C (Effect of Upscaling on Morphometric Accuracy throughout a Microstructural Bone Adaptation Simulation) and D (Effect of *In Vivo* Image Artefacts on Convergence of Upscaled HR-pQCT Simulation). Top: Regularized input models from experiment B were taken and microstructural bone adaptation simulations were run for all input models. Bottom: A separate HR-pQCT dataset was also converted to regularized input models and simulations were run for all input models. For experiment C, static parameters and dynamic parameters were computed. The goal of experiment C was to compare how accurate low-resolution versus upscaled resolution simulations were relative to the reference simulations. For experiment D, static parameters were computed as well. The goal of experiment D was to compare the voxel size dependency of down-scaled micro-CT and rescaled HR-pQCT images to determine if additional artefacts introduced by the HR-pQCT images had a strong influence on the outcome of the bone-adaptation simulation

#### Experiment D: Effect of *In Vivo* Image Artefacts on Convergence of Upscaled HR-pQCT Simulation

Microstructural bone adaptation simulations were run on regularized input models of the *in vivo* HR-pQCT images and their rescaled version (Figure 3) using the same simulation parameters and calculating the same static parameters as in experiment C. The reference BV/TV for the regularized input model generation for all resolutions was based on the respective original HR-pQCT resolution binary structure.

### Evaluation and Statistics

All simulations and evaluations were performed within the trabecular mask. SED was evaluated for non-empty voxels. SED distributions were represented using the SciPy Gaussian kernel density approximation (Virtanen et al. 2019). NumPy (Van Der Walt et al. 2011) was used to compute the Kolmogorov-Smirnov (KS) statistic, mean and standard deviations, as well as BV/TV, which was computed through integration of the model density within the trabecular mask. All other static parameters (Tb.N, Tb.Th, Tb.Sp, SMI) were computed using the scanner manufacturer’s image processing language (IPL) (Hildebrand et al. 1999). Before calling the IPL functions, models were upscaled to 25 μm to remove the voxel-size dependency of IPL functions as a confounding factor. Finally, formed and resorbed volume over time was computed by integration of positive and negative density changes using NumPy.

Comparisons for experiment A were done using a paired Student t-test. For experiments B and C, to determine significance, two-way analysis of variance (two-way ANOVA) was performed for each measured parameter as an omnibus test with the two categorical groups: resolution and upscaling. If heteroscedasticity was detected using a Levene test, heteroscedasticity consistent covariance matrices of type HC3 were used. Post-hoc group comparisons were done using paired Student t-tests and p-values were corrected using the Holm-Bonferroni correction for multiple comparisons. For experiment D, paired Student t-tests were performed and p-values were corrected using the Holm-Bonferroni correction for multiple comparisons. The level of significance was set to 0.05. For the Student t-tests and Levene tests, scipy 1.3.1 was used. The ANOVA was done using statsmodels (Seabold and Perktold 2010) 0.10.2.

## Results

### Experiment A: Effect of Regularized Input Models on Initialisation Shock

The usage of regularized input models removed initialization shocks found in simulations run on conventional input models. For the conventional binary input models, the change in apparent compressive stiffness after the first iteration was a factor of 5.9±0.8 larger than the maximum of all other iteration steps (Figure 4). For BV/TV, a factor of 2.3±1.6 increase was observed in the first iteration (Figure 4). Visually, we observe that for some samples a clear shock in BV/TV was present. In contrast, for the first iteration of the regularized input model approach, the change in apparent compressive stiffness is indistinguishable from the rest of the simulation with a computed factor of 0.5±0.5 increase, which is significantly lower than for the conventional method (p<0.001). For BV/TV, we obtained a factor of 2.1±0.6 increase (Figure 4) which is not significantly different from the threshold method. Since the regularized input model approach removed the initialization shock in apparent stiffness across all samples, we performed all other experiments using this approach.

**Fig 4.**
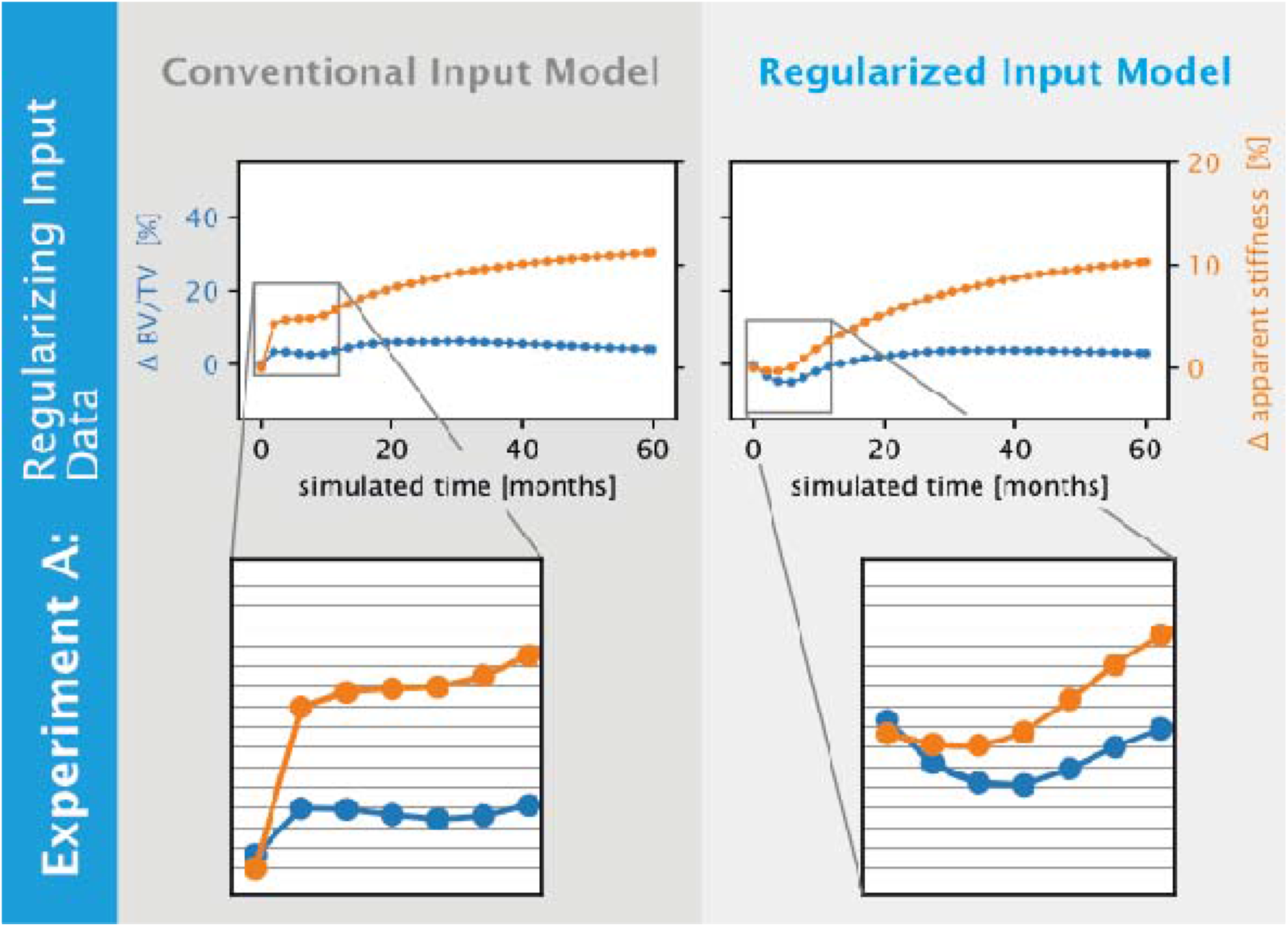
Initialisation shocks in bone volume fraction (BV/TV) and relative apparent stiffness are reduced for regularized input models in comparison to conventional input models, shown for a representative sample of experiment A (Effect of Regularized Input Models on Initialisation Shock). The initialization shock visible in BV/TV and stiffness (left) for the conventional model generation was not present when using the regularization approach (right)

The average regularization threshold for the high-resolution images was 563.6±6.6 mg HA / cm^3^. For the low-resolution images, the average regularization thresholds were 601.8±10.0, 605.2±14.7, and 590.6±18.9 mg HA/ cm^3^ for 40, 61, and 82 μm, respectively.

### Experiment B: Effect of Upscaling on Mechanical and Morphometric Accuracy

For all tested thresholds, the SED distributions for the low-resolution regularized input models did not visually match the SED distribution of the reference regularized input models (Figure 5a). However, for BV/TV matched thresholds, the peaks of the SED distributions aligned well. Qualitatively, the difference in SED distribution from the reference model increased with voxel-size (Figure 5b and 5c). The SED distributions for upscaled regularized input models were almost an order of magnitude closer to the reference than for the low-resolution regularized input models (p<0.05 for all; Table 1). Specifically, deviations in mean SED for upscaled resolutions were less than 5% while deviations were up to 42% for low-resolutions and the KS statistic was below 0.03 for all upscaled resolutions compared to up to 0.22 for low-resolutions (Table 1).

**Fig 5.**
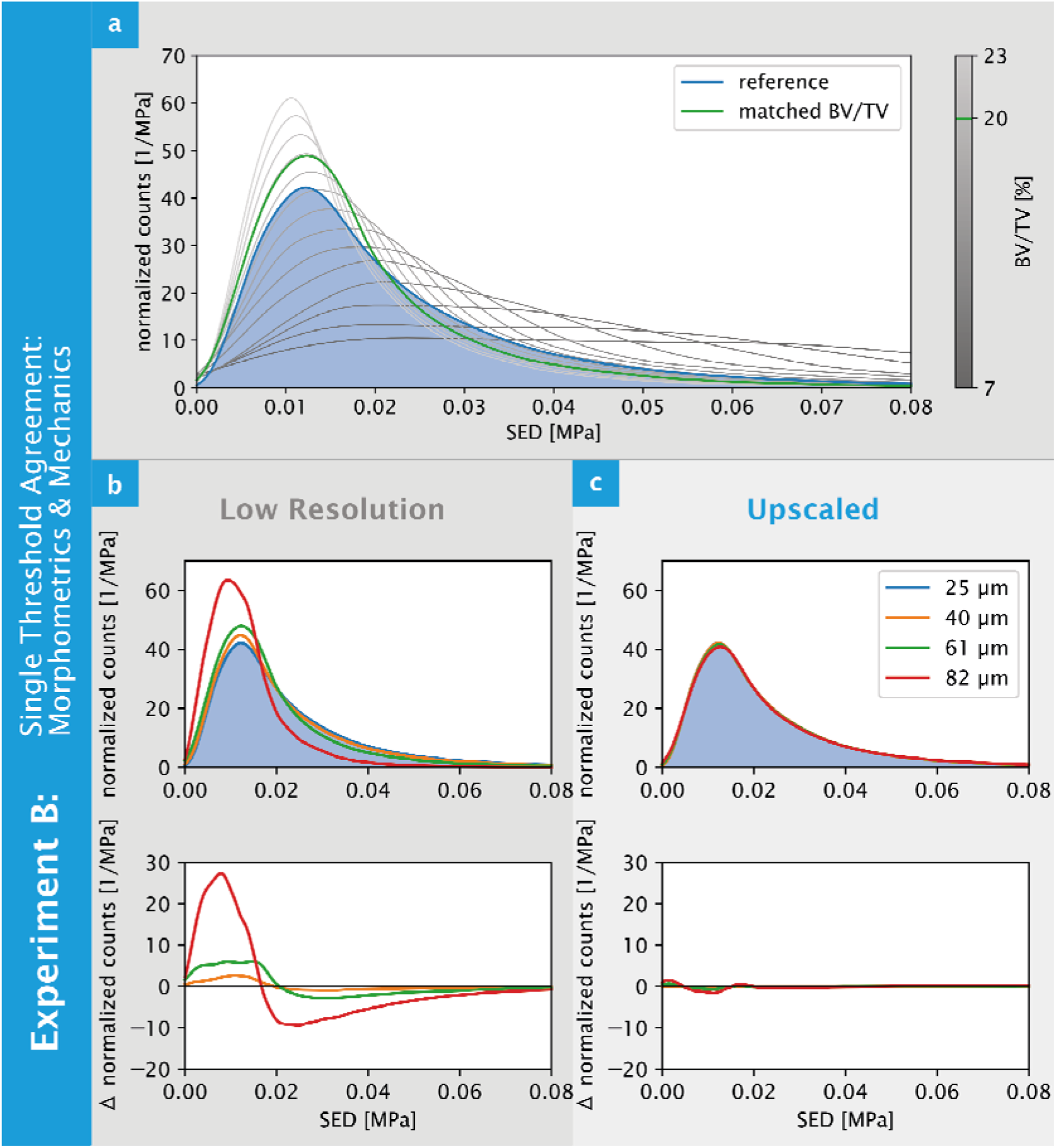
Visual agreement in strain energy density (SED) distributions are improved for upscaled bone volume fraction (BV/TV) matched regularized input models for a representative sample of experiment B (Effect of Upscaling on Mechanical and Morphometric Accuracy). (a) The strain energy density (SED) distributions for the reference resolution (25 μm) (blue) was flatted compared to the down-scaled 61 μm model (green), which used a threshold chosen to match the bone volume fraction (BV/TV) of the reference resolution model. Thresholds corresponding to BV/TV values between 7 and 23% are shown in different shades of grey with a worse agreement with the reference SED distribution. (b)Comparing across all resolutions, the low-resolution models did not capture the SED distribution of the reference model. (c) However, upscaled resolutions clearly captured the distribution of the reference model across all resolutions, with deviations being an order of magnitude smaller compared to models generated without upscaling

**Table 1.**
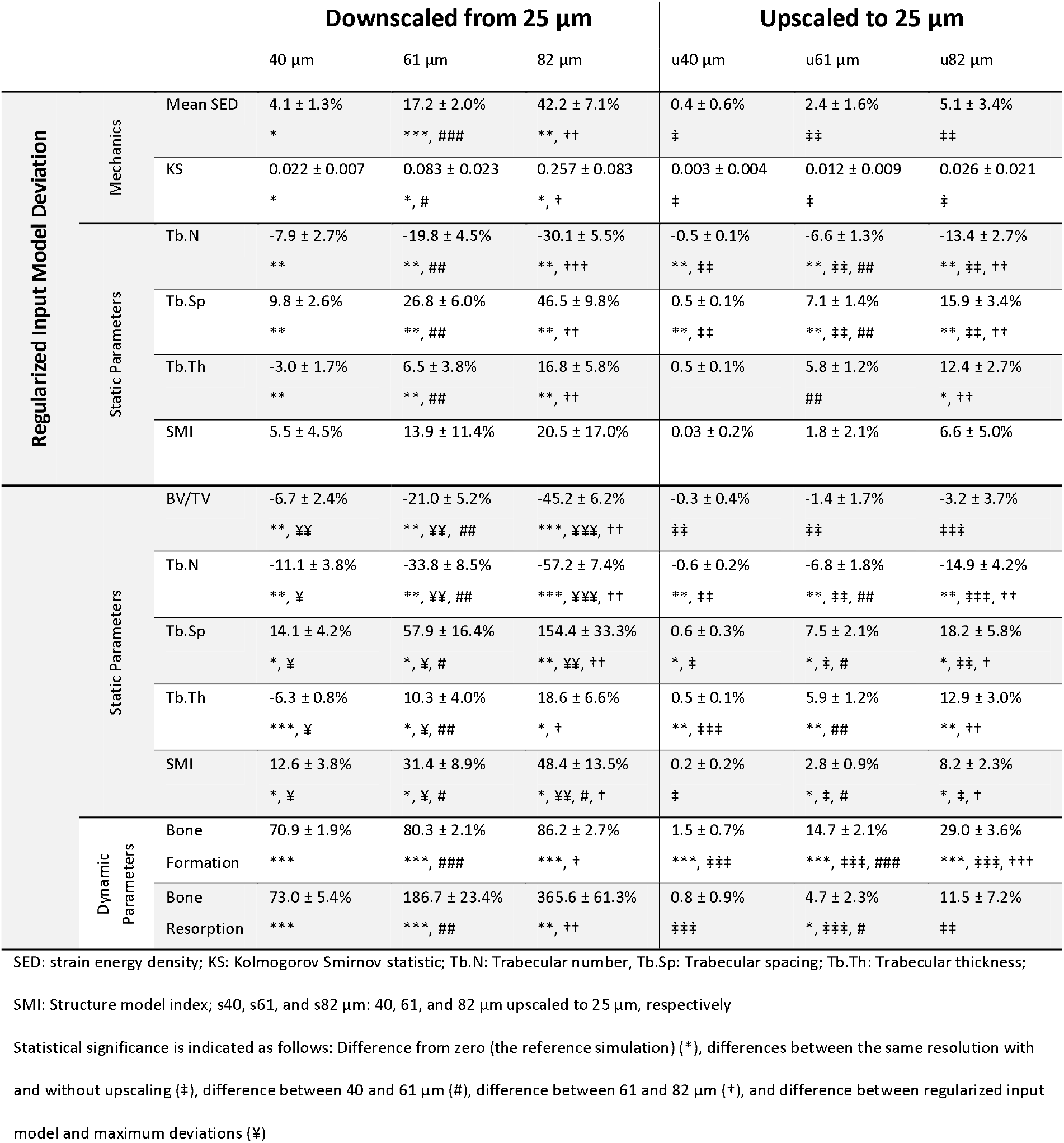
Comparison of microstructural bone adaptation simulation outcomes for regularized model inputs of micro-CT images that were downscaled from low-resolution and then upscaled back to high-resolution. Parameters were compared against the high-resolution micro-CT image reference simulation and relative deviations in percent are shown

Resolution, applied upscaling, and the interaction between resolution and upscaling had a significant effect on all measured static morphometric parameters, except for the interaction of the effects for SMI and the effect of upscaling on Tb.Th (Table 1). Without upscaling, significant differences in the mechanical and static parameters (except SMI) were observed for all lower resolutions (40, 61, and 82 μm) (Table 1). No differences in BV/TV were observed, as BV/TV was matched. With upscaling, deviations in the static parameters were significantly lower for each of the lower resolutions (u40, u61, and u82 μm, respectively) (Table 1).

The average regularization threshold for the upscaled images were 561.7±6.8, 530.9±9.6, and 496.6±13.7 mg HA / cm^3^ for the three upscaled resolutions (u40, u61, and u82 μm), respectively.

### Experiment C: Effect of Upscaling on Morphometric Accuracy Throughout a Microstructural Bone Adaptation Simulation

For the reference simulations, BV/TV was initially reduced in the range of 1.8% to 14.8%, followed by an increase in BV/TV in the range of 3.8% to 38.8% (Figure S1). Tb.N decreased in the range of 5.2% to 22.8%. Trabecular thickness increased in the range of 29.1% to 42.4%, except for one sample for which it decreased by 8.8%. SMI was dependent on the specific sample, with differences in the range of −15.0% to 18.5%; two samples experienced an increase and three samples a decrease in SMI.

Resolution, upscaling, and their interaction had a significant effect (p<0.001) on all measured static and dynamic parameters for the maximum deviations observed during the simulation.

#### Low-Resolution Simulations

Differences in bone-structure between the reference simulation and the low-resolution simulation were visible and increased over the course of the simulation (Figure 6). Deviations in static parameters significantly increased over the course of the simulation compared to the initial models (**Error! Reference source not found**., Figure 7). Compared to the reference simulations, BV/TV and Tb.N were underestimated for all low-resolution simulations (p<0.01), SMI and Tb.Sp were overestimated (p<0.05), and Tb.Th did not follow a clear trend (Figure 77).

**Fig 6.**
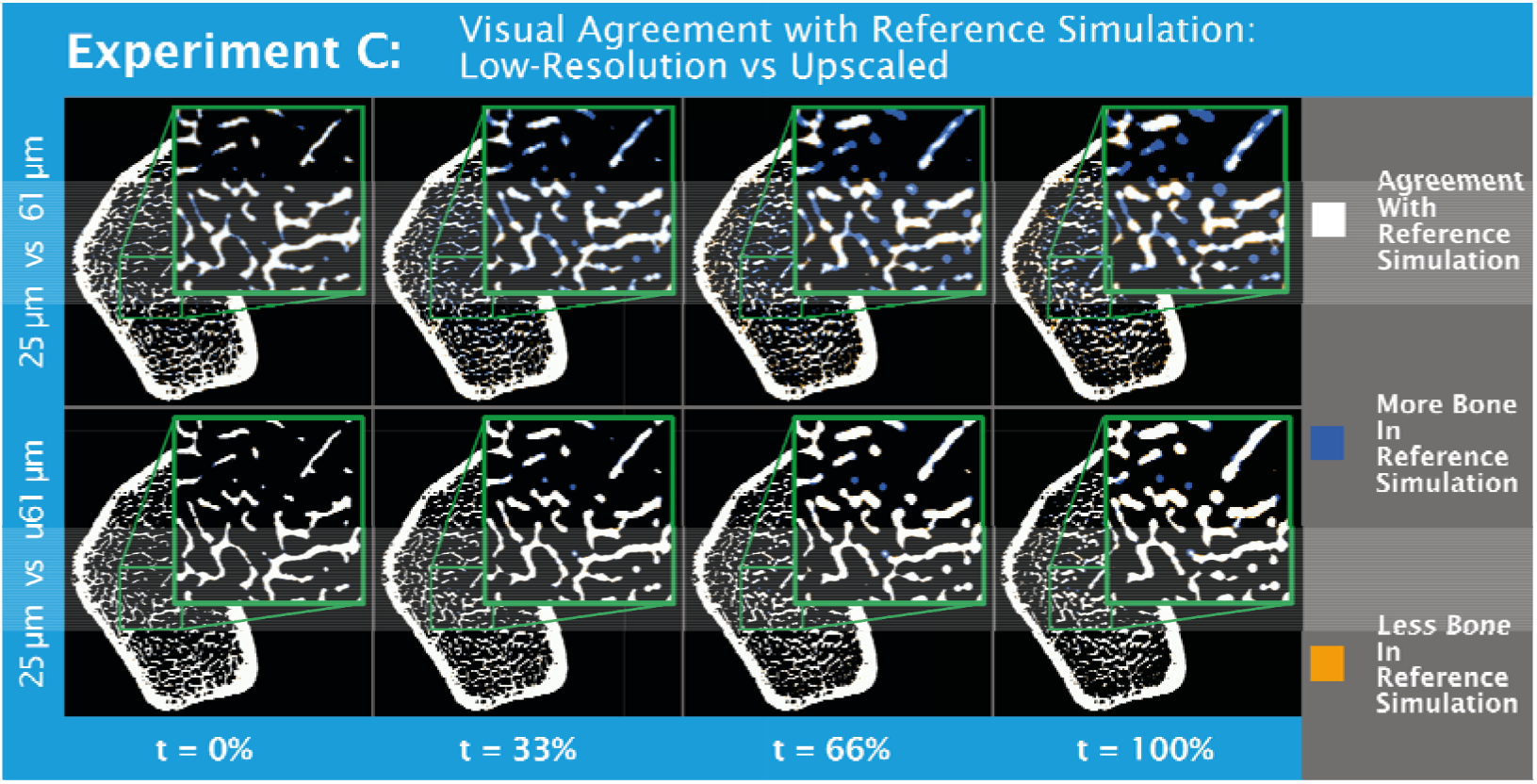
Visual difference after 0, 33, 66, and 100% of the simulated time are reduced for upscaled regularized input models for a representative sample of experiment C (Effect of Upscaling on Morphometric Accuracy Throughout a Microstructural Bone Adaptation Simulation). Top: comparison of reference to low-resolution simulation. Bottom: comparison of reference to upscaled simulation. For the upscaled simulation, very small structures were still lost, due to very thin trabeculae that cannot be captured in a 61 μm image, but the major part of the bone structure remodelled identical to the reference simulation for upscaled images

**Fig 7.**
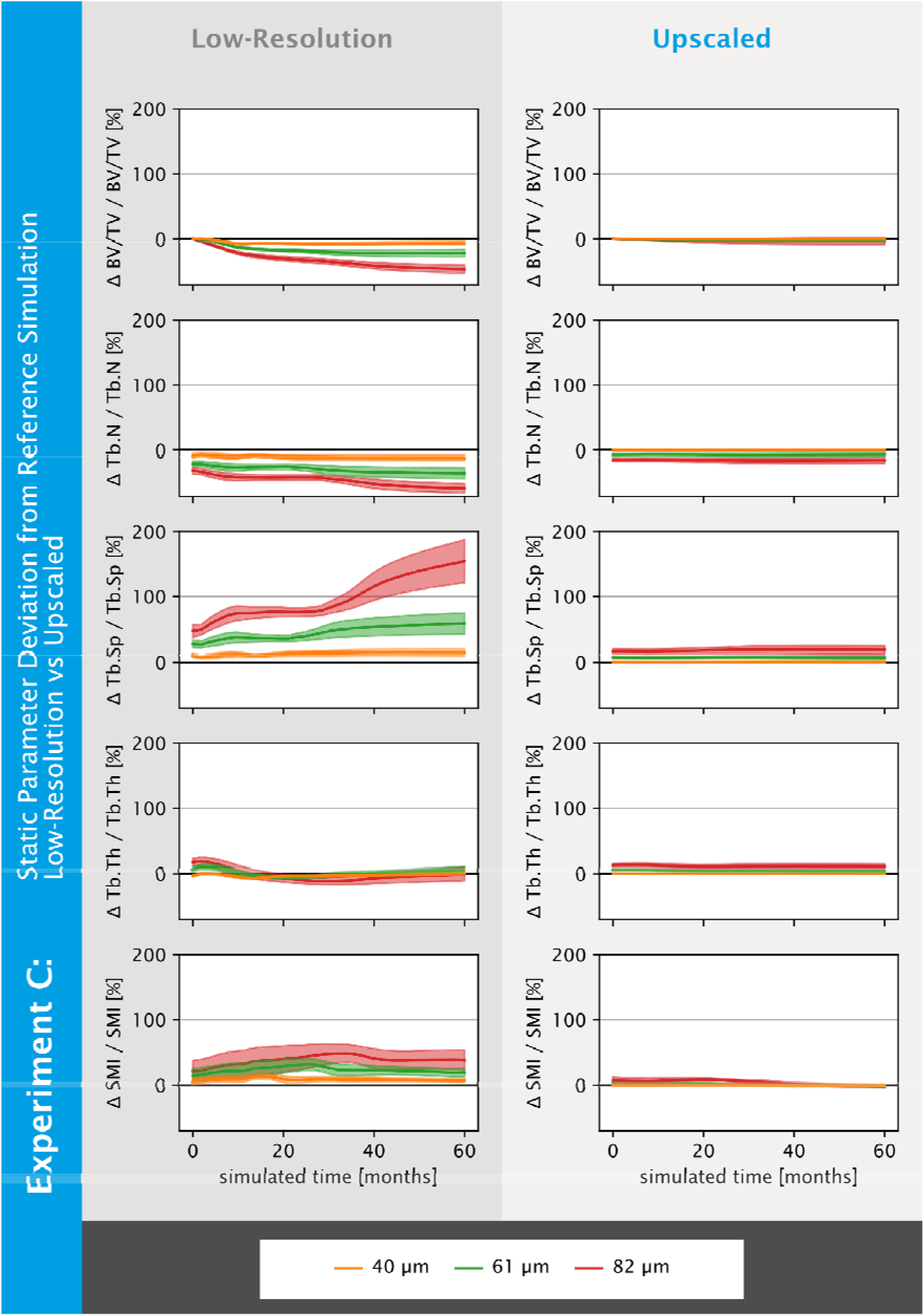
Deviations from the reference simulation in static parameters are reduced for upscaled regularized input models across all resolutions as found in experiment C (Effect of Upscaling on Morphometric Accuracy Throughout a Microstructural Bone Adaptation Simulation). Deviations in static parameters from the reference simulation (25 μm) over the course of the simulation for low-resolution (left) and upscaled regularized input models (right) are shown. Computed static parameters are: Bone volume fraction (BV/TV), trabecular number (Tb.N), trabecular spacing (Tb.Sp), trabecular thickness (Tb.Th), and structure model index (SMI). BV/TV was matched for the initial model, resulting in perfect agreement between the resolutions. Generally, for the different samples, upscaling improves the agreement with the reference (25 μm) simulations. Deviations were more predictable after upscaling of the image

Bone formation and resorption rates were significantly larger for low-resolution simulations compared to the reference simulations (Table 1). Visually, the formation rate for the low-resolution simulations peaked at a later time point and had lower peak values compared to the reference simulation (Figure 8). For the resorption rate, peak delay and widening was also observed for the low-resolution simulations (Figure 8). However, the magnitude of the resorption rate peaks increased with voxel-size and the initial resorption rate decayed slower compared to formation rates, for which the opposite was observed.

**Fig 8.**
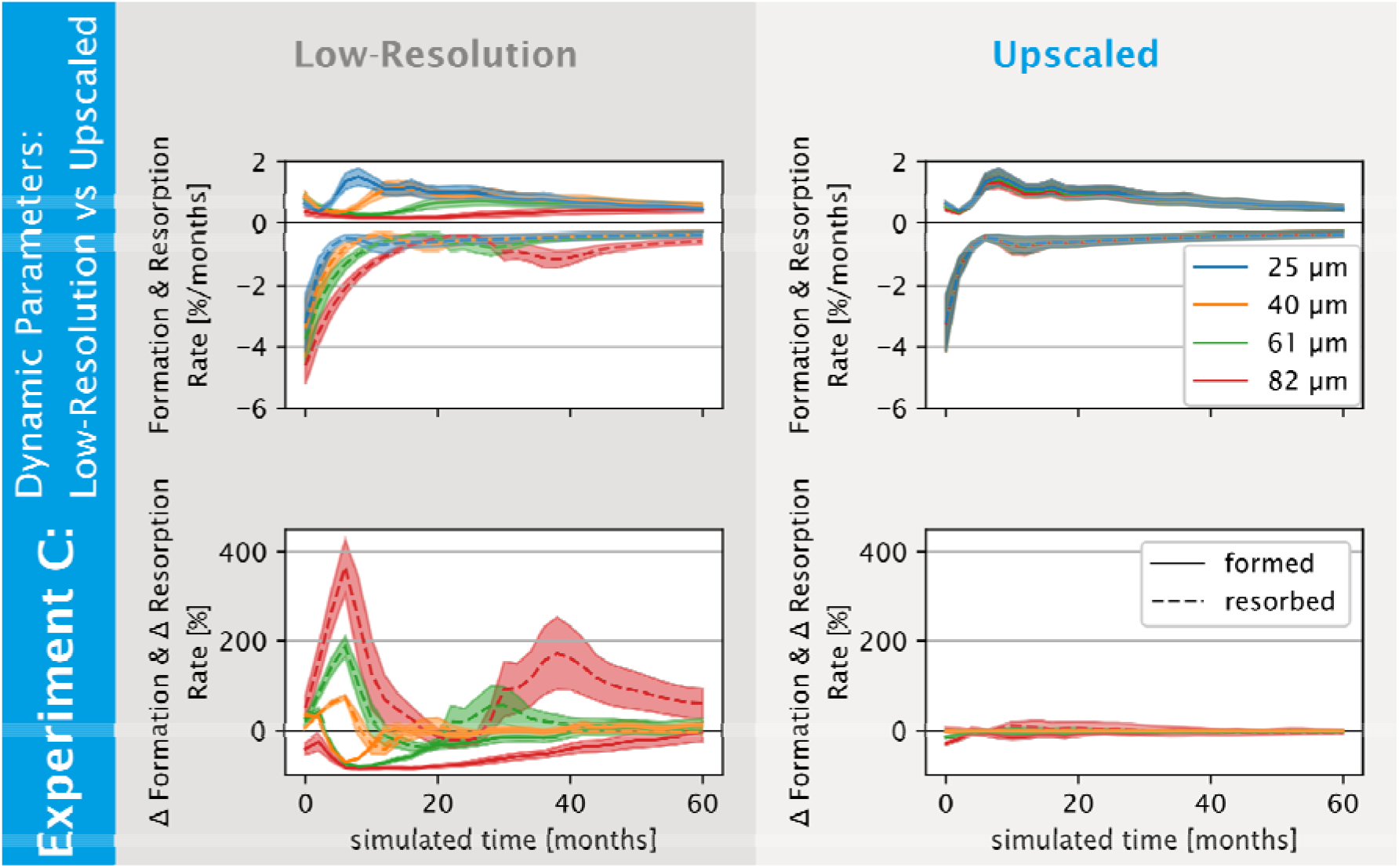
Deviations from the reference simulation in dynamic parameters are reduced for upscaled regularized input models across all resolutions as found in experiment C (Effect of Upscaling on Morphometric Accuracy Throughout a Microstructural Bone Adaptation Simulation). Dynamic parameter results of simulation run on low-resolution (left) and upscaled regularized input models (right) are shown. Bone formation and resorption over time (top), and deviations of these parameters from the reference simulation (25 μm) (bottom). Reference simulation results were aligned with the use of upscaled regularized input models with respect to the amount and time-point of formation and resorption events. Deviations were an order of magnitude smaller for the simulations run on upscaled regularized input models

**Fig 9.**
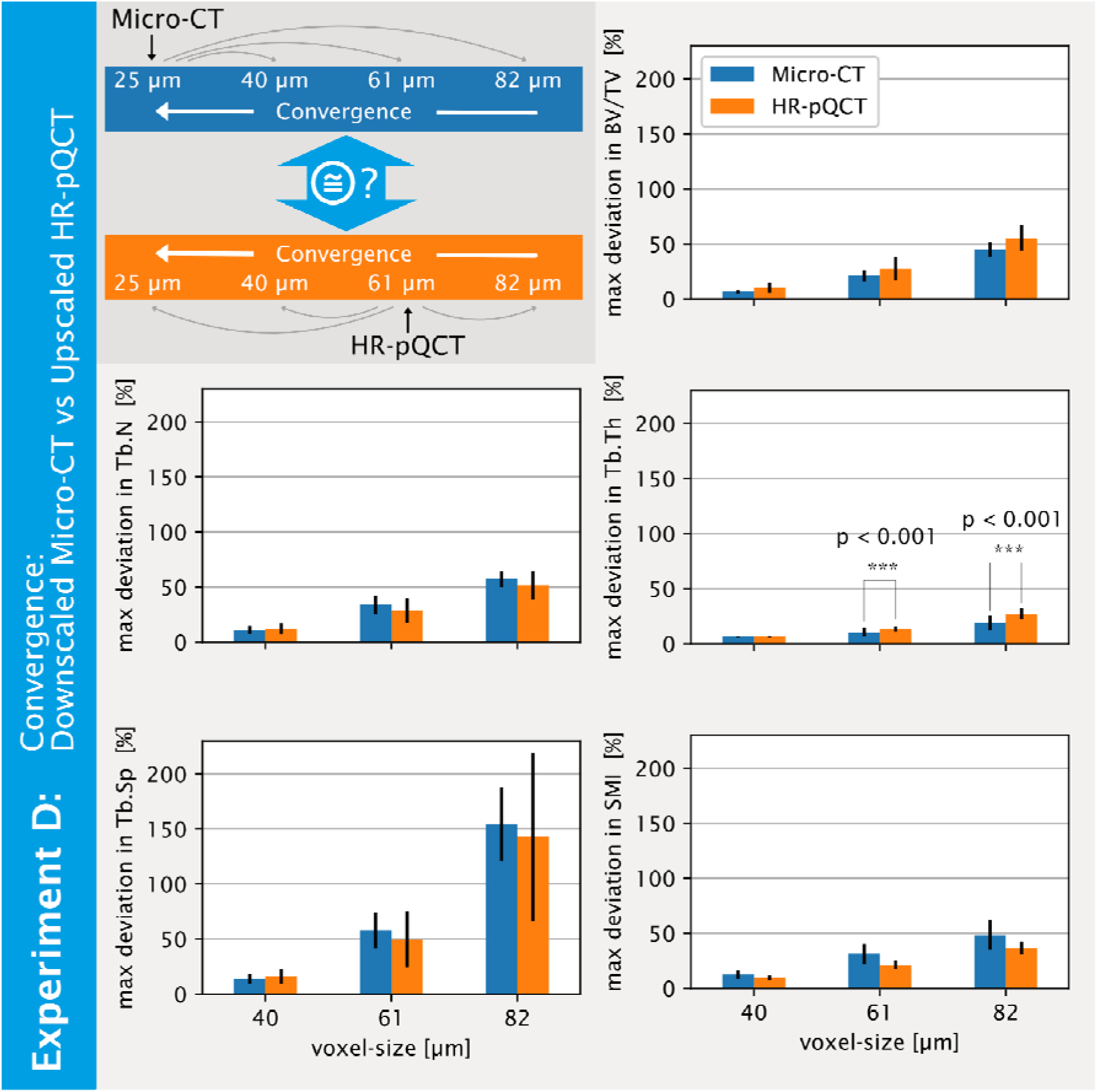
Agreement in convergence between regularized input models from downscaled micro-CT and rescaled HR-pQCT as found in experiment D (Effect of *In Vivo* Image Artefacts on Convergence of Upscaled HR-pQCT Simulation). Convergence behaviour of low-resolution simulations from experiment C (Effect of Upscaling on Morphometric Accuracy Throughout a Microstructural Bone Adaptation Simulation) and from rescaled HR-pQCT image simulations (top left). Maximum mean deviations from the reference (25 μm) simulations and corresponding standard deviations are shown for all computed static parameters. Significant differences were observed for trabecular thickness (Tb.Th); the mean maximum deviations of all other parameters are not significantly different, indicating that voxel-size was the dominating factor on the simulation outcomes

#### Upscaled Simulations

Visually, differences in bone structure over the course of the simulation compared to the reference simulations were drastically reduced for the upscaled simulations (Figure 6). Accuracy in BV/TV was improved by an order of magnitude for the highest available clinical resolution (s61 μm) compared to the low-resolution simulations. The accuracy of all other static parameters was also significantly improved (Table 1, Figure 77). The maximum deviations in static parameters were not significantly greater than those of the input models for all upscaled resolutions.

For the dynamic parameters, accuracy of bone formation and resorption per time unit was significantly increased for the upscaled images (Table 1). The formation and resorption rates peaked at the same time-point across all resolutions, within the temporal resolution of the simulation. The peaks of both rates were of similar magnitude across all resolutions. Visually, the overall shapes of the formation and resorption curves were similar across all resolutions for the duration of the simulation period, with no noticeable widening or shift of peaks (Figure 88).

### Experiment D: Effect of *In Vivo* Image Artefacts on Convergence of Upscaled HR-pQCT Simulations

After an initial drop in BV/TV in the range of 7.2% to 24.2%, the simulations on the *in vivo* HR-pQCT data upscaled to 25μm showed varying behaviour. Three samples showed an increase in BV/TV, one sample had a close to stable BV/TV over time, and one sample experienced a further reduction in BV/TV before BV/TV began increasing after half of the simulation time. Tb.N. decreased for all samples over time in the range of 12.0% to 42.9%, whereas Tb.Th. increased over time in the range of 13.8% to 65.1%. Tb.Sp also increased in the range of 13.6% to 78.0%. SMI decreased for all samples in the range of 7.8% to 21.8%. The spread of SMI values across all samples decreased over the course of the simulation by 47.8%.

Comparing the convergence of the different static parameters with respect to resolution between the low-resolution simulations from experiment C and the simulations from experiment D, no significant differences could be found except for Tb.Th at 61 and 82 μm resolutions (p<0.001) (Figure 99). Hence, the effects of noise and other additional imaging artefacts from *in vivo* HR-pQCT were smaller than the effects of model resolution, which dominated the convergence errors observed in the static parameters (Figure 99).

## Discussion

The objectives of this study were to investigate whether microstructural simulations of *in vivo* human HR-pQCT images yielded accurate results and were a viable tool as part of a VPH.

### Experiment A: Effect of Regularized Input Models on Initialisation Shock

Since one of the ideas of the VPH is to provide doctors with a decision support system (Viceconti and Hunter 2016), the goal of every microstructural bone adaptation simulation must be to achieve parity between the simulated structure and the structure observed *in vivo*. We observed that when using conventional model inputs, the apparent compressive stiffness showed an initialization shock behaviour (Figure 4), with a change approximately six times larger than any other change in stiffness over the course of the simulation. While initialisation shocks have not been studied in the context of microstructural bone adaptation simulations, in the context of ocean climate models, Mulholland et al. (Mulholland et al. 2015) identified that the removal of certain model components can result in abrupt changes in the dynamics of the system. The analogy for microstructural bone adaptation simulations are the mismatch of applied boundary conditions and the true, but unknown, *in vivo* boundary conditions. This mismatch can also be interpreted as the removal of certain boundary condition forces at the beginning of the *in silico* adaptation. We tackled this challenge by employing the load estimation algorithm by Christen et al. (2013) which tries to estimate the *in vivo* applied loads more closely than the uniaxial compression boundary conditions typically used with H R-pQCT radius data (Burghardt et al. 2011).

Another potential cause for initialisation shocks could be the abrupt change of surface geometry from *in vivo* microstructural bone adaptation to *in silico* bone adaptation simulations. Using a regularized input, we used information in the grey-scale image that is normally cut off, improving the input model generation to reduce the initialisation shock and the associated effect on the results. For all simulations, the developed regularized input model approach removed the shock behaviour in apparent compressive stiffness (Figure 4). While the magnitude of the change in BV/TV for the initial iteration step did not change with the new regularized input model, change in BV/TV looks smoother using this approach (Figure 4). Importantly, larger changes in BV/TV are expected for this type of simulation, as the structure adapts to the applied boundary conditions over the course of the simulation, yielding less changes as the structure reaches a shaped optimized to the applied boundary conditions. We conclude that the first iteration step does not have to be excluded if this new approach is used, which allows direct comparisons of morphometrics and mechanics to *in vivo* measurements. Furthermore, the fact that the initialisation shock was removed might indicate that the regularized input model is a more accurate mechanical representation of the *in vivo* bone structure than the conventional binary version.

### Experiment B: Effect of Upscaling on Mechanical and Morphometric Accuracy

Another obstacle to overcome when running microstructural bone adaptation simulations on *in vivo* HR-pQCT images was finding an accurate digital representation of a bone captured *in vivo* with HR-pQCT with respect to mechanics and morphometry (Alsayednoor et al. 2018), which is an obvious requirement of bone adaptation simulations. We found that with the use of upscaling, the choice of a single threshold provided regularized input models that agreed well for BV/TV, mechanical properties, and other tested morphometric parameters (i.e. Tb.N, Tb.Sp, and SMI). Furthermore, the linear trend of the thresholds for the different upscaled resolutions indicated that even in the absence of a high resolution ground truth, an appropriate threshold for accurate morphometrics and mechanics can be chosen. In contrast, our results for images that were not upscaled agreed with previous research (Alsayednoor et al. 2018), which showed no agreement between mechanics and morphometrics for various thresholds (Figure 5). This lack of agreement also holds true for thresholds optimized to match BV/TV (Figure 5), a method used in a previous study by Christen et al. to investigate the voxel size dependence of a micro-FE based load estimation algorithm (Christen et al. 2016). The culmination of these results indicates that upscaling may also be useful for other applications using images of HR-pQCT resolution.

### Experiment C: Effect of Upscaling on Morphometric Accuracy Throughout a Microstructural Bone Adaptation Simulation

To study the accuracy of microstructural bone adaptation simulations for images with HR-pQCT resolutions, we used high-resolution micro-CT images as ground truth, as this method has been previously validated to simulate realistic bone structures over time (Badilatti et al. 2016). The use of different low-resolution voxel-sizes (40, 61, and 82 μm) resulted in deviations in morphometric parameters of more than 30% in comparison to the reference simulations, however deviations as small as 15% may indicate disease, such as osteoporosis (Zhang et al. 2010). Importantly, observed differences in parameters were not consistent, thus could not be corrected or recovered via calibration curves, as has been possible in previous studies (Müller et al. 1996). In comparison to static morphologic parameters, dynamic parameters were more dependent on image resolution. This can be understood by realizing that static parameters are averaged over the bone structure, such that local differences in morphometrics, e.g. trabecular number, may be concealed. On the other hand, dynamic parameters are additive and sum all changes over the given volume. Thus, even minor differences in structure result in larger deviations in dynamic parameters. Importantly, standard deviations of dynamic parameters from a previous study (Schulte et al. 2013) were lower by up to an order of magnitude compared to the deviations we observed for the highest clinically available resolution (80% vs 38% for formation, 187% vs 18% for resorption) (Table 1). Therefore, the accuracy we observed would be insufficient given the magnitude of natural variation previously observed for these dynamic parameters. Overall, our results show that microstructural bone adaptation simulations run on native clinical scanner resolutions suffer from poor accuracy in the assessment of static morphometric and dynamic parameters; thus, limiting future use as a model for human bone adaptation.

Running the same simulations on the upscaled images (u40, u61, and u82 μm) resulted in a drastic reduction in static parameter deviations to less than 10%. For BV/TV, these deviations were near 1% (Table 1), which is similar to the reproducibility limit of BV/TV for the clinical setting (0.84-1.14%) (MacNeil and Boyd 2008; Mueller et al. 2009). The improved accuracy in assessment of static parameters from the upscaled images is likely due to the need for higher resolution to correctly represent the reference model in terms of mechanics and morphology (Figure 6, Table 1). Any deviations of the regularized input model led to error accumulation throughout the simulation, such as missing thin individual trabecular structures that grow in thickness in the reference simulations (Figure 6). These errors ultimately led to larger morphometric deviations of the final structure (Figure 77). Only the dynamic parameters obtained from the upscaled simulations correctly captured the overall curve profile of the high-resolution simulations, matching the position, width, and height of peak formation and resorption rates. The large deviations of the downscaled resolutions indicate that there may have also been an intrinsic voxel-size dependence of the algorithm, independent of the initial model. The observed maximum deviations in dynamic parameters are also at least a factor of two smaller than the natural variation observed in inbred mice indicating sufficient accuracy of the method for human *in vivo* applications (Schulte et al. 2013).

### Experiment D: Effect of *In Vivo* Image Artefacts on Convergence of Upscaled HR-pQCT Simulations

Finally, we investigated the effects of clinically observed image artefacts, e.g. higher noise levels, by using *in vivo* HR-pQCT images. Comparing the simulations run on these images to those run on upscaled versions of the same images, no significant difference was observed in the convergence of the different static parameters (Figure 99), except for Tb.Th. Since the *ex vivo* and *in vivo* dataset are not from the same study participants, it is possible that this difference in Tb.Th is due to unknown physiological differences between the *in vivo* subject group and the *ex vivo* samples. Furthermore, with respect to the overall deviation observed in Tb.Th, the observed significant difference is still small, especially given the fact that Tb.Th is known to be difficult to capture with HR-pQCT resolution (MacNeil and Boyd 2007; Manske et al. 2015) and are therefore likely not clinically relevant.

### Limitations

This study is, however, not without limitations. One limitation of this study was the lack of a high resolution ground truth scan of the patient radii used in Experiment D. However, the artefacts most commonly associated with *in vivo* HR-pQCT images, such as motion artefact, are difficult to recreate with cadaveric specimen, while micro-CT images cannot be obtained from patients due to the radiation dosage and imaging volume. Thus, we utilized both high-resolution cadaveric images and clinically acquired *in vivo* HR-pQCT images of patients to assess these factors independently.

An additional limitation of this study is the sample size (n=5 for both *ex vivo* and *in vivo* experiments). However, the small spread in deviations across subjects observed from the results of the upscaled simulations indicates that a larger sample size may not be warranted. Importantly, the inclusion of additional samples would have required an excess of computational resources due to the high resolution of the simulations.

## Conclusions

In conclusion, we found model resolution to be the dominating image property which drove convergence errors in microstructural bone adaptation simulations. Importantly, upscaling drastically reduced errors in mechanical analyses, static morphological parameters, and dynamic parameters, resulting in simulation outcomes that, even for clinically available resolutions, were similar to those from high-resolution images. Initialisation errors were avoided with the use of upscaling and the proposed regularization method, which generated model input that closely represented the true bone structure with respect to both mechanics and morphometry. With these results, we conclude that microstructural bone adaptation simulations can be run on *in vivo* HR-pQCT images and yield realistic results, given a validated set of parameters. Hence, these simulations provide a powerful tool to study disease related bone microstructure changes in patients, as part of the VPH vision.

## Acknowledgements

Funding for the DACH Fx Project from the Swiss National Science Foundation (Lead Agency, 320030L_170205), German Research Foundation (IG 18/19-1, SI 2196/2-1), and Austrian Science Fund (I 3258-B27) is gratefully acknowledged. This work was supported by a grant from the Swiss National Supercomputing Centre (CSCS) under project ID s841.

## Supplementary Figure S1

**Fig S1.**
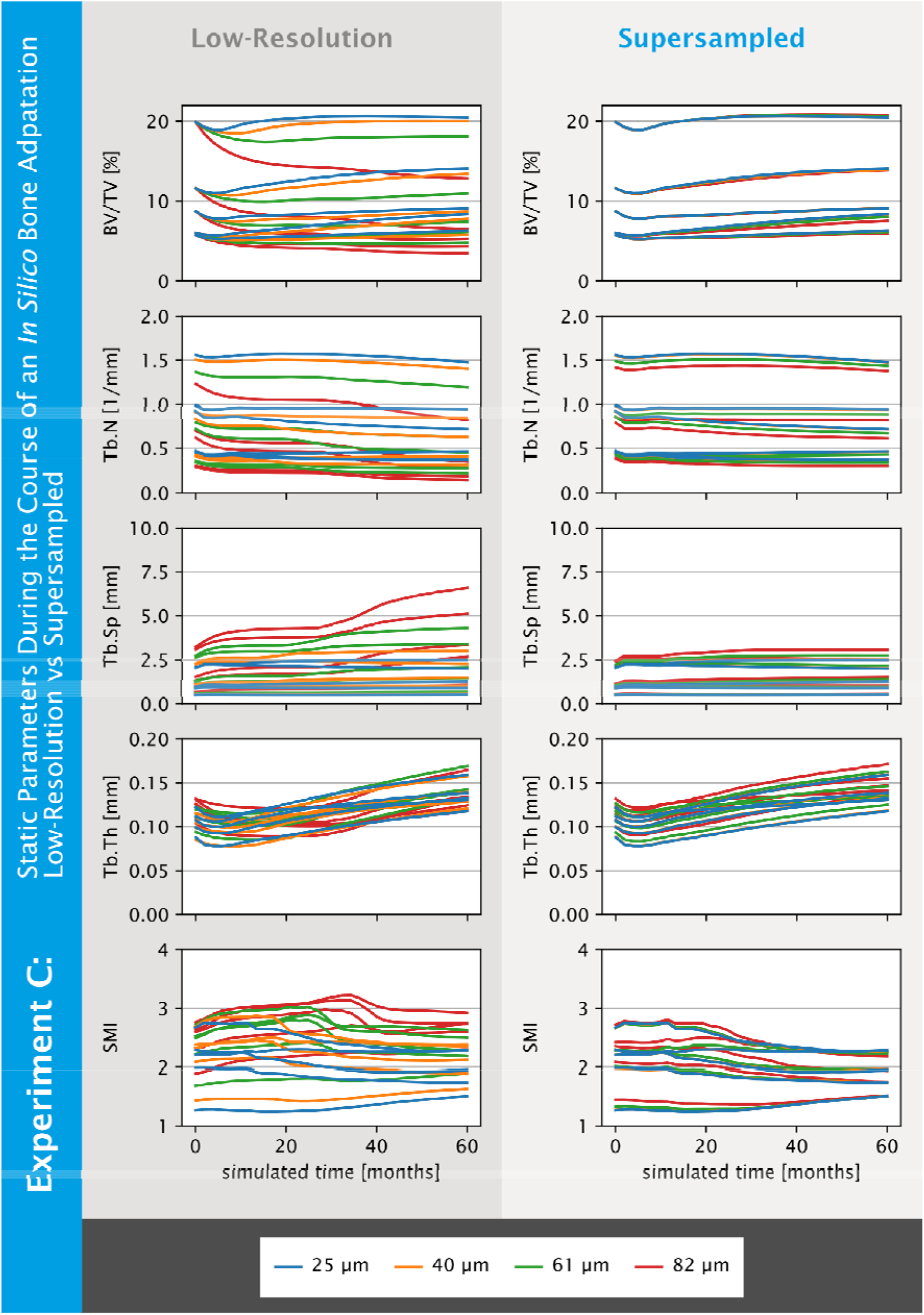
Visual deviations from the reference simulation in static parameters are reduced for upscaled regularized input models across all resolutions as found in experiment C (Effect of Upscaling on Morphometric Accuracy Throughout a Microstructural Bone Adaptation Simulation). Static parameters over the course of the simulation for low-resolution (left) and upscaled regularized input models (right) are shown. Computed static parameters are: Bone volume fraction (BV/TV), trabecular number (Tb.N), trabecular spacing (Tb.Sp), trabecular thickness (Tb.Th), and structure model index (SMI). BV/TV was matched for the initial model, which is why initially we got perfect agreement between the resolutions. Overall we see for the different samples that upscaling improves the agreement with the reference (25 μm) simulations

